# High-quality nuclear genome and mitogenome of *Bipolaris sorokiniana* strain LK93, a devastating pathogen causing wheat root rot

**DOI:** 10.1101/2022.12.28.522073

**Authors:** Wanying Zhang, Qun Yang, Lei Yang, Haiyang Li, Wenqing Zhou, Jiaxing Meng, Yanfeng Hu, Limin Wang, Ruijiao Kang, Honglian Li, Shengli Ding, Guotian Li

## Abstract

*Bipolaris sorokiniana*, one of the most devastating hemibiotrophic fungal pathogens, causes root rot, crown rot, leaf blotching, and black embryos of gramineous crops worldwide, posing a serious threat to global food security. However, the host-pathogen interaction mechanism between *B. sorokiniana* and wheat remains poorly understood. To facilitate related studies, we sequenced and assembled the genome of *B. sorokiniana* strain LK93. Nanopore long reads and next generation sequencing short reads were applied in the genome assembly, and the final 36.4 Mb genome assembly contains 16 contigs with the contig N50 of 2.3 Mb. Subsequently, we annotated 11,811 protein-coding genes including 10,620 functional genes, 258 of which were identified as secretory proteins including 211 predicted effectors. Additionally, the 111,581 bp mitogenome of LK93 was assembled and annotated. The LK93 genomes presented in this study will facilitate research in the *B. sorokiniana*-wheat pathosystem for better control of crop diseases.

## Genome Announcement

The devastating fungal pathogen *Bipolaris sorokiniana* causes a variety of cereal crop diseases like root rot and black point of wheat (DOLAR et al., 2019; Al-Sadi, 2021; Sharma et al., 2021), resulting in significant declines in grain yield and quality (Fatemeh et al., 2022). The diseases such as root rot and black point of wheat commonly developed in winter wheat of the Huanghuai floodplain, one of the major wheat production areas in China, which produces two-thirds of wheat grains in China (Zhao, 2010). Through the identification of wheat black point pathogen in this region, a total of 41 varieties were susceptible to *B. sorokiniana* among the 50 tested wheat varieties (Dai et al., 2011). Similarly, *B. sorokiniana* was isolated from winter wheat with rotten roots in this region (Kang et al., 2020). Although genome assemblies of *B. sorokiniana* from barley and wheat have been reported (Condon et al., 2013; Darshan et al., 2020; Meng et al., 2020; Rashmi et al., 2022), the high-quality genome for *B. sorokiniana* strains isolated from the Huanghuai floodplain was lacking. Previous studies suggested that *B. sorokiniana* varied a lot among different geographical populations (Gurung et al., 2013; Kang et al., 2020). Isolated from the Huanghuai floodplain of China, LK93 shows strong pathogenicity (Ma et al., 2022; Zhang et al., 2022), and is considered as a suitable material to study pathogenicity and test resistance in different wheat cultivars to *B. sorokiniana*. Therefore, we sequenced and assembled a high-quality genome of a *B. sorokiniana* strain isolated from this region of China (Zhang et al., 2022).

We isolated 673 isolates from diseased wheat samples collected from 97 locations in the Huanghuai floodplain from 2014 to 2015, among which 262 isolates were identified as *B. sorokiniana* (Kang et al., 2020). Out of these isolates, *B. sorokiniana* strain LK93 normal in conidiation and vegetative growth-isolated from Lankao, China-was selected for genome sequencing (Ma et al., 2022; Zhang et al., 2022).

High-quality genomic DNA was extracted from 48-h-old mycelia, and the methods of fungal culture and DNA extraction were referred to Wang (Wang et al., 2022). We amplificated and sequenced the internally transcribed spacer (ITS) region, and the sequence comparison results showed that the ITS region of LK93 is 100% identical to *B. sorokiniana* (accession MH208982.1). The high-quality genomic DNA sequencing was performed on the PromethION platform of the Oxford Nanopore technology and on the MGISEQ-2000RS platform of the next generation sequencing (NGS) technology, respectively, at Nextomics Biosciences (Wuhan, China). Consequently, a total of 7.4 Gb Nanopore long reads, and 4.4 Gb NGS short reads were obtained. To guarantee that only high-quality long reads were used in the assembly, the long reads were filtered out removing reads shorter than one kb or with a Phred quality score less than seven. Based on filtered long reads, the LK93 genome was *de novo* assembled with NextDenovo (v2.5.0). Likewise, the short reads were filtered out to eliminate reads with the q value smaller than 20. To improve the assembly quality, three rounds of base correction with filtered short and long reads were performed using NextPolish (v1.3.1) (Hu et al., 2020) based on the draft genome. The final genome assembly is 36.4 Mb, consisting of 18 contigs (N50 = 2.3 Mb). Analyzed by D-genes (Cabanettes and Klopp, 2018), high collinearity was found between the assembly and the reference genomes (BRIP27492a, WAI2406) (McDonald et al., 2019) (Fig. S1-S2). To obtain a chromosome-level genome, the assembly was anchored into 16 scaffolds of the reference genome BRIP27492a via the RAGOO (v1.11) (Alonge et al., 2019) scaffolding method. Two unanchored short contigs (159,427 bp) that considered to be mitochondrial fragments were deleted. Telomere sequences of 16 scaffolds were tested and integrated into the Table S1. Additionally, using RepeatMasker (v4.1.0) (Tarailo-Graovac and Chen, 2009) and RepeatModeler (v2.0.1) (Flynn et al., 2020), we identified and masked 5.3 Mb repetitive sequences in the LK93 nuclear genome. The final genome assembly, evaluated by Benchmarking Universal Single-Copy Orthologs (BUSCO) (v4.0.1) (Simão et al., 2015) based on the fungi_odb10 lineage dataset, contains 99.3% of the complete ortholog groups, showing the high-quality of the LK93 genome. The assembly statistics are summarized in Table 1. GC content and gene density of the genome assembly were analyzed in nonoverlapping 100-kb intervals (Chen et al., 2020; Chen et al., 2022) (Fig. 1).

**Table 1.**
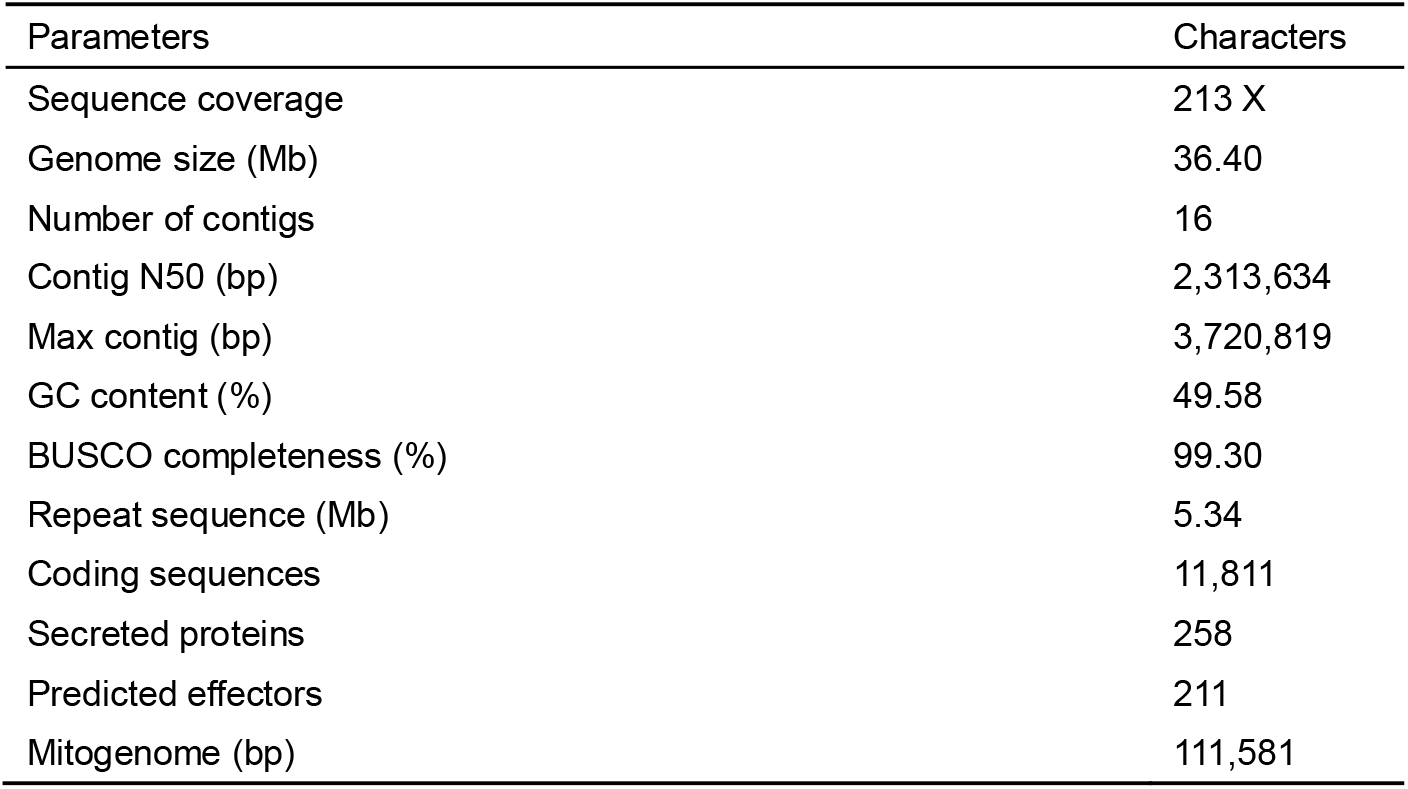
Summary of the *B. sorokiniana* LK93 genome

**Fig. 1.**
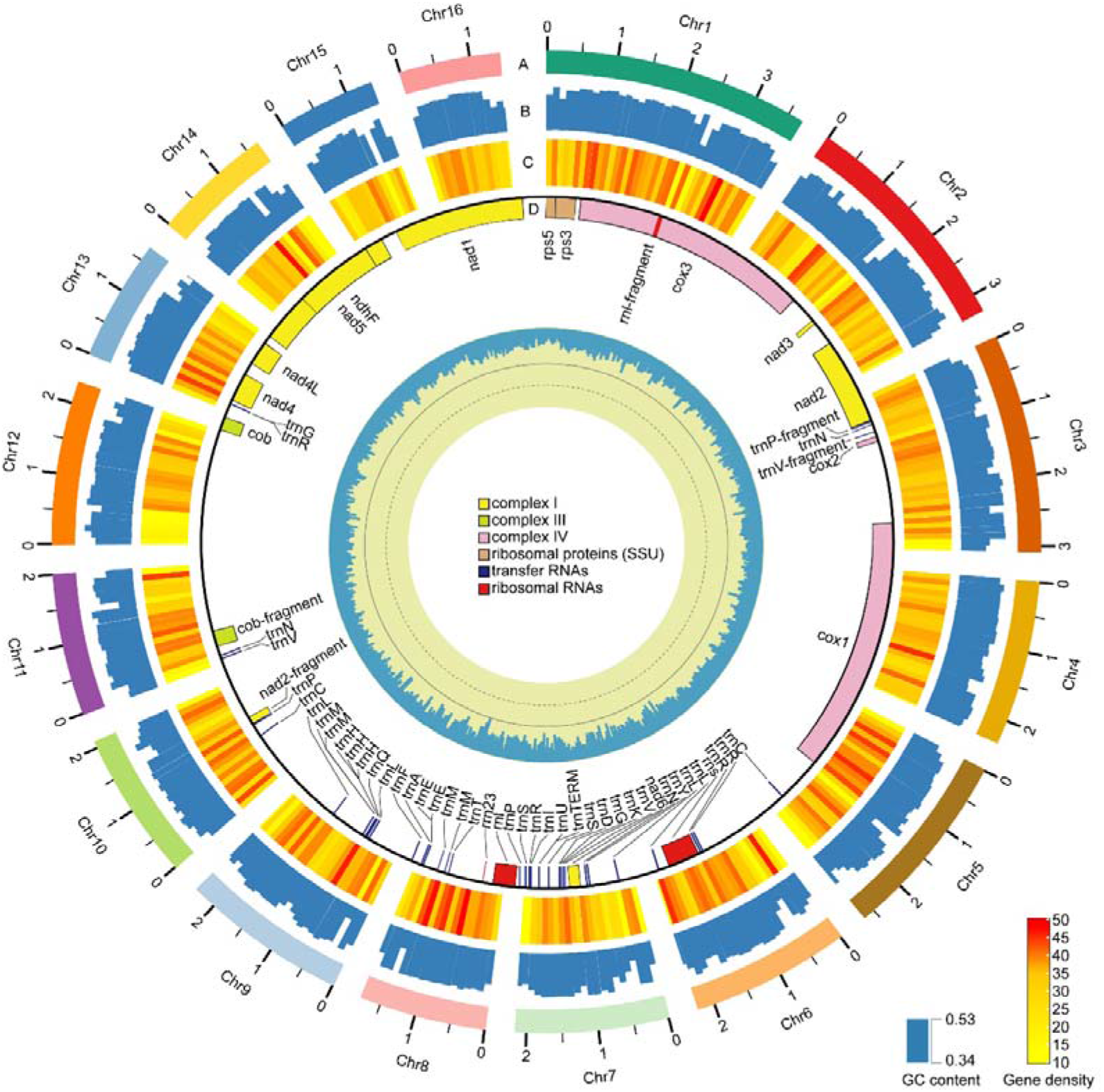
Nuclear genome and mitogenome of the *B. sorokiniana* strain LK93. The 16 chromosomes on a megabase scale, GC content, and gene density are visualized in circles A, B, C, respectively. Data in circles B and C are displayed in nonoverlapping 100-kb intervals. The mitogenome and predicted genes are displayed in circle D. The innermost circle shows the mitogenome in the kb scale as well as the GC content. Tracks A, B and C share the same scale.

For gene annotation of the *B. sorokiniana* LK93 genome, a total of 9.0 Gb RNA-seq data were obtained from mycelia and infected wheat roots at different stages (Zhang et al., 2022). The subsequent genome annotation was performed via the Funannotate (v1.8.9) pipeline based on the filtered RNA-seq data. Trinity (v2.8.5) (Grabherr et al., 2011) was used to assemble transcriptome and HISAT (v2.2.1) (Kim et al., 2019) was used to map the RNA-seq reads to the genome in the annotation progress. Augustus (v3.3.3) (Stanke et al., 2006) and PASA (v2.4.1) were used for gene prediction and annotation. We annotated 11,811 protein-coding sequences, including 1,191 hypothetical proteins, and 10,620 functional proteins in the LK93 genome (Table 1). Taking advantage of SignalP (v5.0) (Almagro Armenteros et al., 2019), TMHMM (v2.0) (Möller and Croning, 2002), and EffectorP (v3.0) (Sperschneider and Dodds, 2021), we annotated 258 secretory proteins in the genome, and 211 of which were identified as candidate effectors including 112 cytoplasmic effectors and 99 apoplastic effectors that could facilitate plant infection such as CsSp1 (Condon et al., 2013; Zhang et al., 2022).

Fungal mitochondria carry partial genetic information, and the mitogenome plays an important role in evolutionary biology and systematics of fungi (Song et al., 2020). Based on the NGS short reads, we assembled and further annotated the circular whole mitogenome of LK93 assembly with GetOrganelle (v1.7.5.3) (Jin et al., 2020) and Geseq (Michael et al., 2017), respectively. A total of 23,483,390 reads were used in the mtDNA assembly and the slimmed FASTA assembly Graph file that can be visualized by Bandage (Wick et al., 2015) to show the circular path is available in the Figshare website. The length of the LK93 mitogenome is 112 kb. The LK93 mitogenome contains three type complexes, including nine NADH dehydrogenases, two ubiquinol cytochrome c reductases, and three cytochrome c oxidases. Also, we identified an ATP synthase, two ribosomal proteins (*rps3* and *rps5*), and 46 RNAs (42 tRNAs and 4 rRNAs) in the mitogenome, similar to results published by Song and colleagues (Song et al., 2020). The circular mitogenome and relevant annotation information were visualized in Figure 1 utilizing OGDRAW (Stephan et al., 2019).

In our study, we present the high-quality nuclear genome and a complete mitogenome of *B. sorokiniana* strain LK93 isolated from Chinese Huanghuai floodplain. We annotated 11,811 protein-coding genes of which 10,620 are functional. A total of 258 secretory proteins including 211 candidate effectors were predicted in the genome assembly. These resources facilitate studies of pathogenicity of*B. sorokiniana* and provide valuable information for protection of cereal crops.

## Supporting information

Supplemental Software Parameters

## Data Availability

The LK93 genome file is available at NCBI under project accession number PRJNA869783. All of the data generated in this study, including the nuclear genome annotation file, the sequence file of candidate effectors, the mitogenome file, and the mitogenome annotation file have been deposited to Figshare database (https://doi.org/10.6084/m9.figshare.20484390).

