## Supplementary figures and images for "High-quality nuclear genome and mitogenome of *Bipolaris sorokiniana* strain LK93, a devastating pathogen causing wheat root rot"

### Supplemental Figure S1

**Supplemental Figures**

**Figure S1** Dot plot comparing the LK93 and BRIP27492a genomes

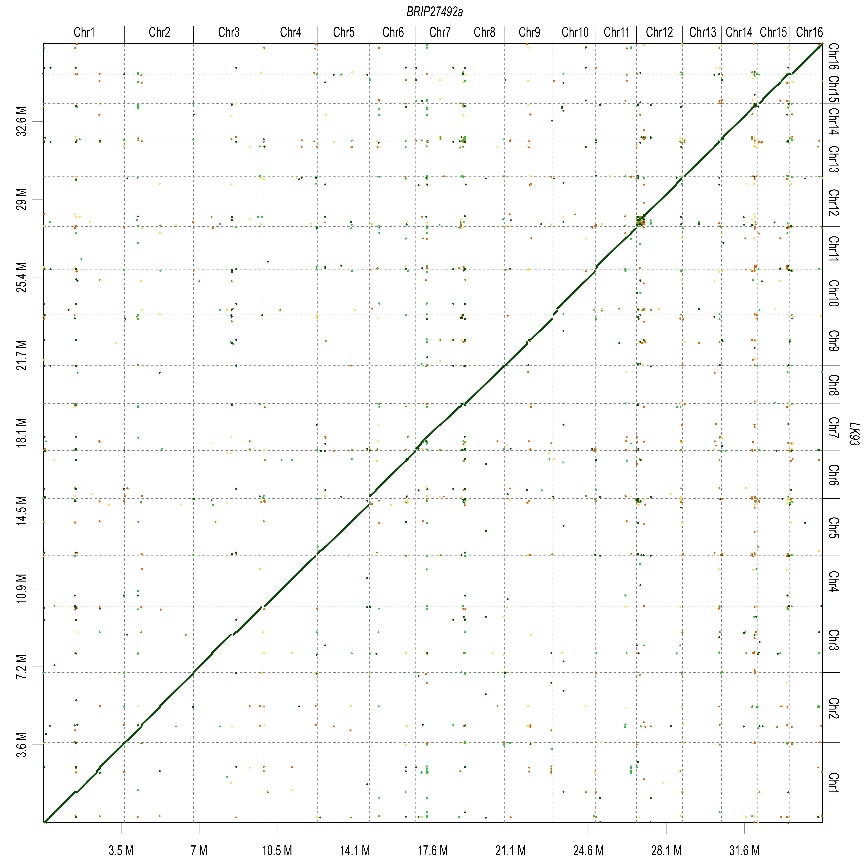

### Supplemental Figure S2

**Supplemental Figures**

**Figure S2** Dot plot comparing the LK93 and WAI2406 genomes
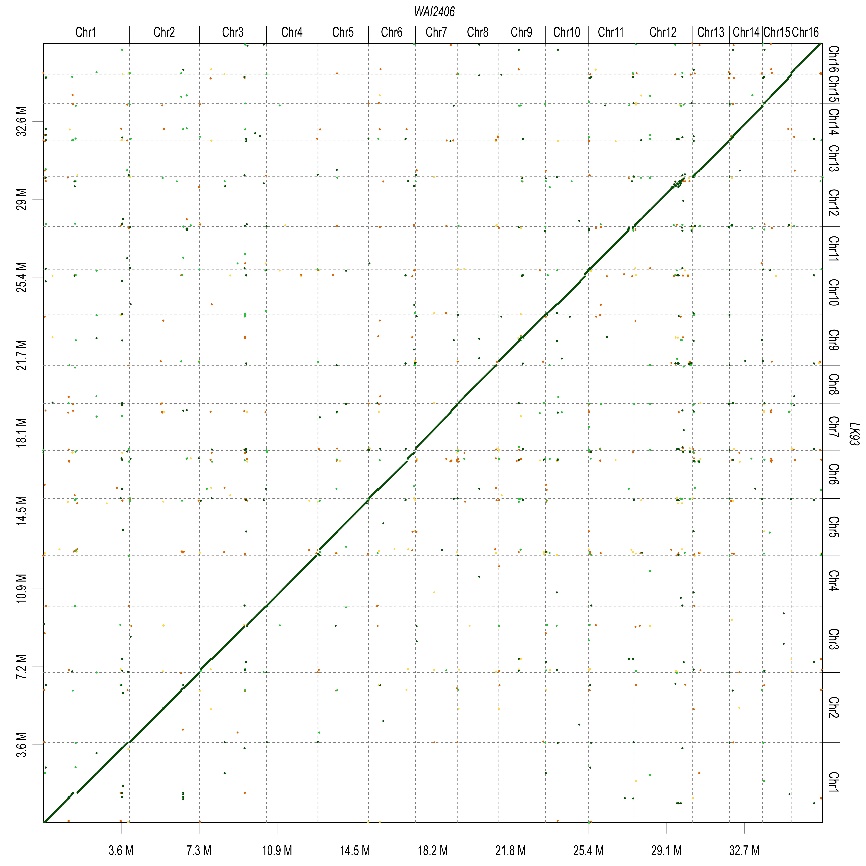
