## Supplemental Table S1 for "High-quality nuclear genome and mitogenome of *Bipolaris sorokiniana* strain LK93, a devastating pathogen causing wheat root rot"

**Supplemental Table S1** Telomere sequences of LK93 chromosomes

| SeqID | SeqLength | Start | End | Length | Type |
| --- | --- | --- | --- | --- | --- |
| Chr2 | 3259185 | 3259048 | 3259083 | 36 | TTAGGG |
| Chr3 | 3091669 | 3091507 | 3091620 | 114 | TTAGGG |
| Chr5 | 2668290 | 69 | 170 | 102 | CCCTAA |
| Chr5 | 2668290 | 2668205 | 2668228 | 24 | TTAGGG |
| Chr6 | 2239657 | 14 | 79 | 66 | CCCTAA |
| Chr7 | 2162158 | 3 | 92 | 90 | CCCTAA |
| Chr8 | 1790119 | 1790002 | 1790103 | 102 | TTAGGG |
| Chr10 | 2095247 | 27 | 92 | 66 | CCCTAA |
| Chr10 | 2095247 | 2095153 | 2095176 | 24 | TTAGGG |
| Chr11 | 2012514 | 11 | 118 | 108 | CCCTAA |
| Chr11 | 2012514 | 2012360 | 2012497 | 138 | TTAGGG |
| Chr12 | 2313634 | 9 | 122 | 114 | CCCTAA |
| Chr13 | 1712079 | 1711967 | 1712068 | 102 | TTAGGG |
| Chr14 | 1686337 | 43 | 78 | 36 | CCCTAA |
| Chr14 | 1686337 | 1686248 | 1686301 | 54 | TTAGGG |
| Chr15 | 1381986 | 11 | 124 | 114 | CCCTAA |
| Chr15 | 1381986 | 1381890 | 1381985 | 96 | TTAGGG |
| Chr16 | 1425224 | 1425047 | 1425166 | 120 | TTAGGG |
