## Supplemental Software Parameters for "High-quality nuclear genome and mitogenome of *Bipolaris sorokiniana* strain LK93, a devastating pathogen causing wheat root rot"

[General]

job_type = local

job_prefix = nextDenovo

task = all

rewrite = yes

deltmp = yes

parallel_jobs = 5

input_type = raw

read_type = ont

input_fofn = lgs.fofn

workdir = assembly

[correct_option]

read_cutoff = 1k

genome_size = 37m

sort_options = -m 20g -t 8

minimap2_options_raw = -t 8

pa_correction = 3

correction_options = -p 8

[assemble_option]

minimap2_options_cns = -t 8

nextgraph_options = -a 1
